## Supplemental Figures for "Presynaptic NMDA receptors cooperate with local action potentials to implement activity-dependent GABA release from the reciprocal olfactory bulb granule cell spine"

**Supplemental material**

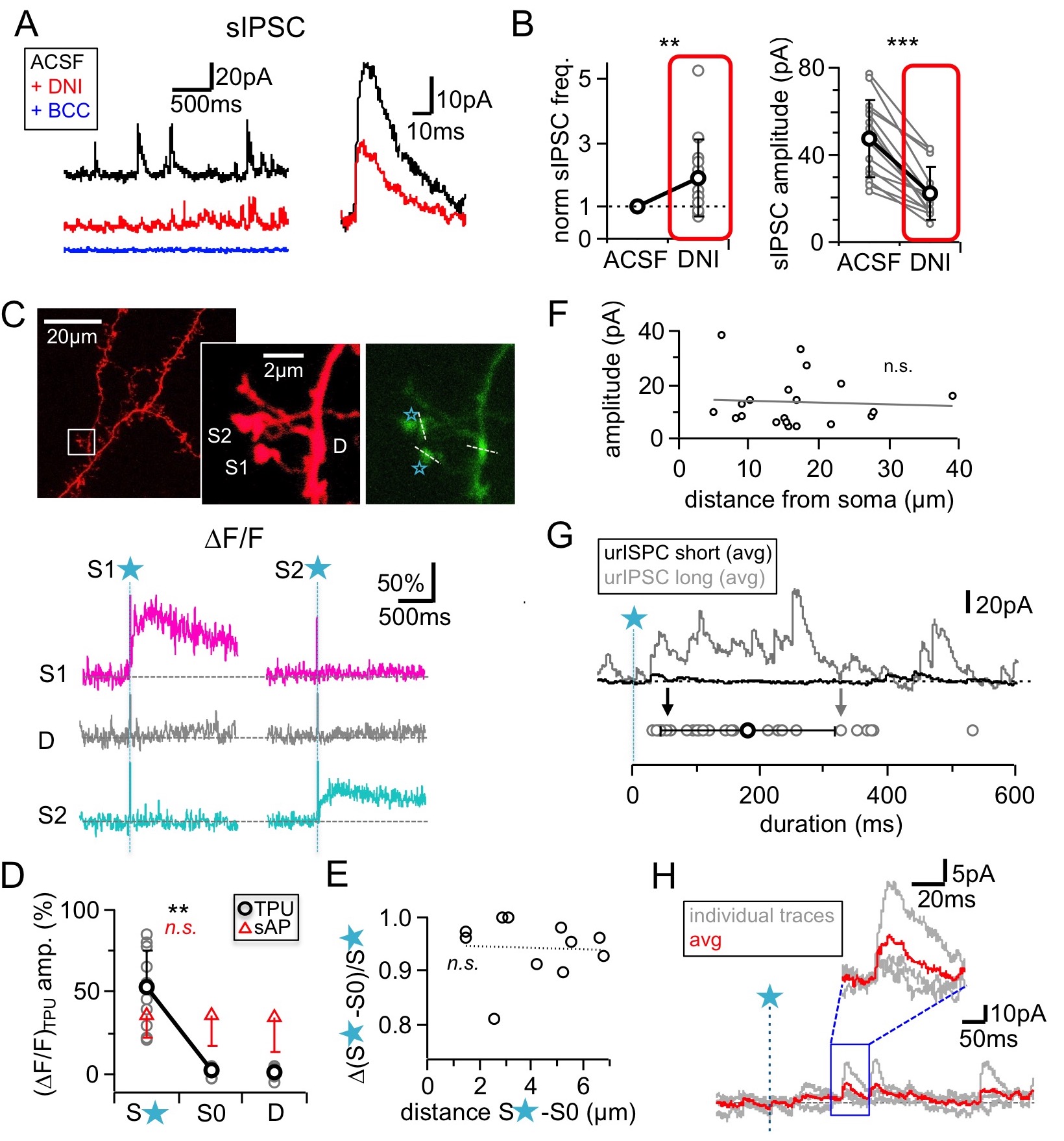

**Figure S1. Effects of DNI (1 mM) on sIPSCs, spine-specificity of TPU, TPU sites, duration of responses**

(**A**) Left: Example recordings in ACSF, after DNI wash-in and with added bicuculline (BCC, 50 µM). Right: Representative single events before and after DNI wash-in.

(**B**) Summary of effects of DNI (red) on spontaneous IPSCs. Left: Frequency. Right: Absolute amplitude. (n = 14 MCs each).

(**C**) Effect of uncaging on non-stimulated adjacent GC spines (S1, S2). Representative experiment. Top: Z-projections of two-photon scan of GC dendrite and spine pair filled with Alexa594 and OGB-1 (right panel, see Methods). Locations of TPU marked with blue stars, location of line scans marked with white dashed lines. Bottom: Averaged fluorescence transients recorded in both spine heads and dendritic shaft in between in response to TPU at S1 only (left) and TPU at S2 only (right).

(**D**) Cumulative data from 11 spine pairs. Grey circles: Individual ∆F/F amplitudes in stimulated spine, non-stimulated spine S0 and dendritic shaft. Black circles: mean values. Red triangles: Mean ∆F/F amplitudes in response to single somatically evoked action potential. No significant difference across all three compartments.

(**E**) Cumulative data from the same 11 spine pairs. Linear regression between relative difference in ∆F/F in the stimulated vs the non-stimulated spine head (y-axis) and the distance between the spine head centers (x-axis). No correlation was found (Pearson’s correlation coefficient r = -0.17, P = 0.31).

(**F**) Distance of TPU site from soma versus urIPSC amplitudes. There was no detectable correlation (n = 20 MCs, Pearson’s correlation coefficient r = -0.07, P = 0.39).

(**G**) Top: Exemplary averaged responses from two experiments. Bottom: Cumulative durations of triggered events (black and grey arrow: durations of exemplary averaged responses).

(**H**) Exemplary experiment with a long latency of first detectable response (see Methods). Inset: Magnification of responses.

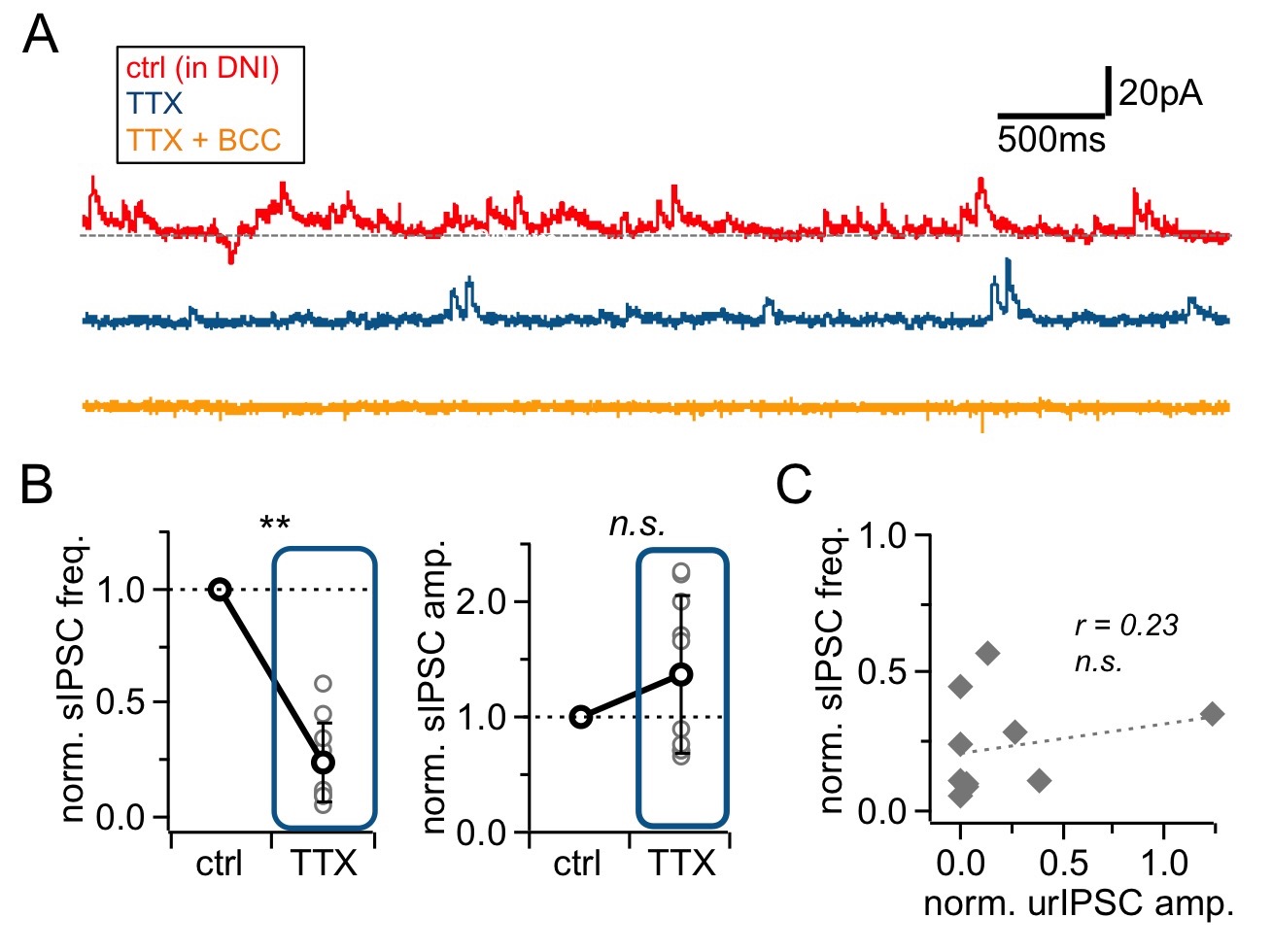

**Figure S2. Effects of Na_v_ blockade (TTX, 500 nM) on spontaneous activity.**

(**A**) Representative experiment showing voltage-clamp recordings (+10 mV) from a mitral cell in the presence of DNI (control, top trace) upon addition of TTX (middle trace) and further wash-in of BCC (50 µM; bottom trace).

(**B**) sIPSCs in control and in presence of TTX (n = 10 MCs). Left: Frequency. Right: Amplitude. sIPSCs were abolished in BCC (analysis not shown) (**C**) Linear regression between the TTX effect on sIPSC frequency versus urIPSC amplitude (n = 10 MCs). No significant correlation.

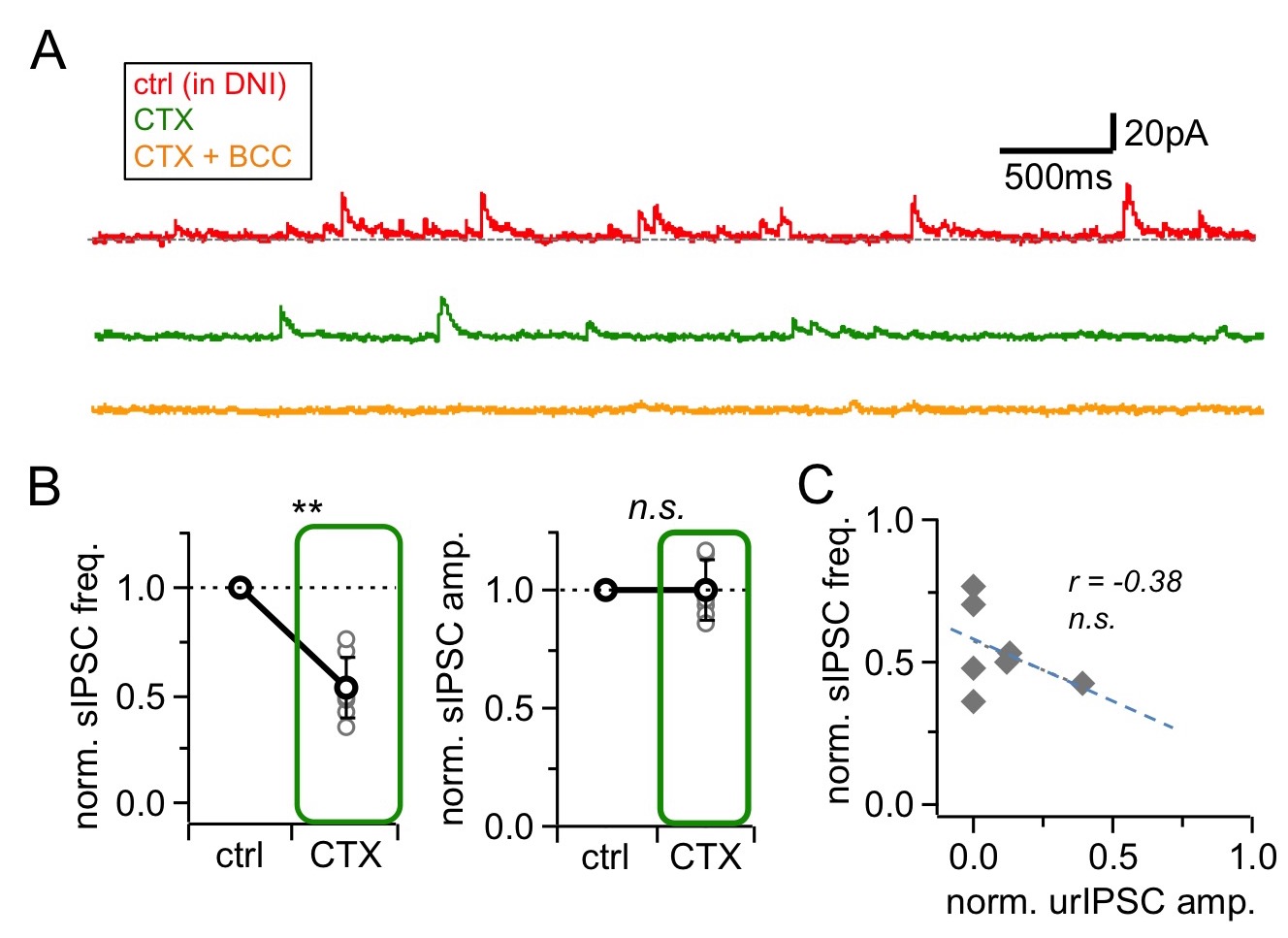

**Figure S3. Effects of HVACC blockade by ω-conotoxin MVIIC (CTX, 1 μM) on spontaneous activity.**

(**A**) Representative experiment showing voltage-clamp recordings (+10 mV) from a mitral cell in the presence of DNI (control, top trace) upon addition of CTX (middle trace) and further wash-in of BCC (50 µM; bottom trace).

(**B**) sIPSCs in control and in presence of CTX (n = 7 MCs). Left: Frequency. Right: Amplitude. sIPSCs were abolished in BCC (analysis not shown) (**C**) Linear regression between the CTX effect on sIPSC frequency versus urIPSC amplitude (n = 7 MCs). No significant correlation.

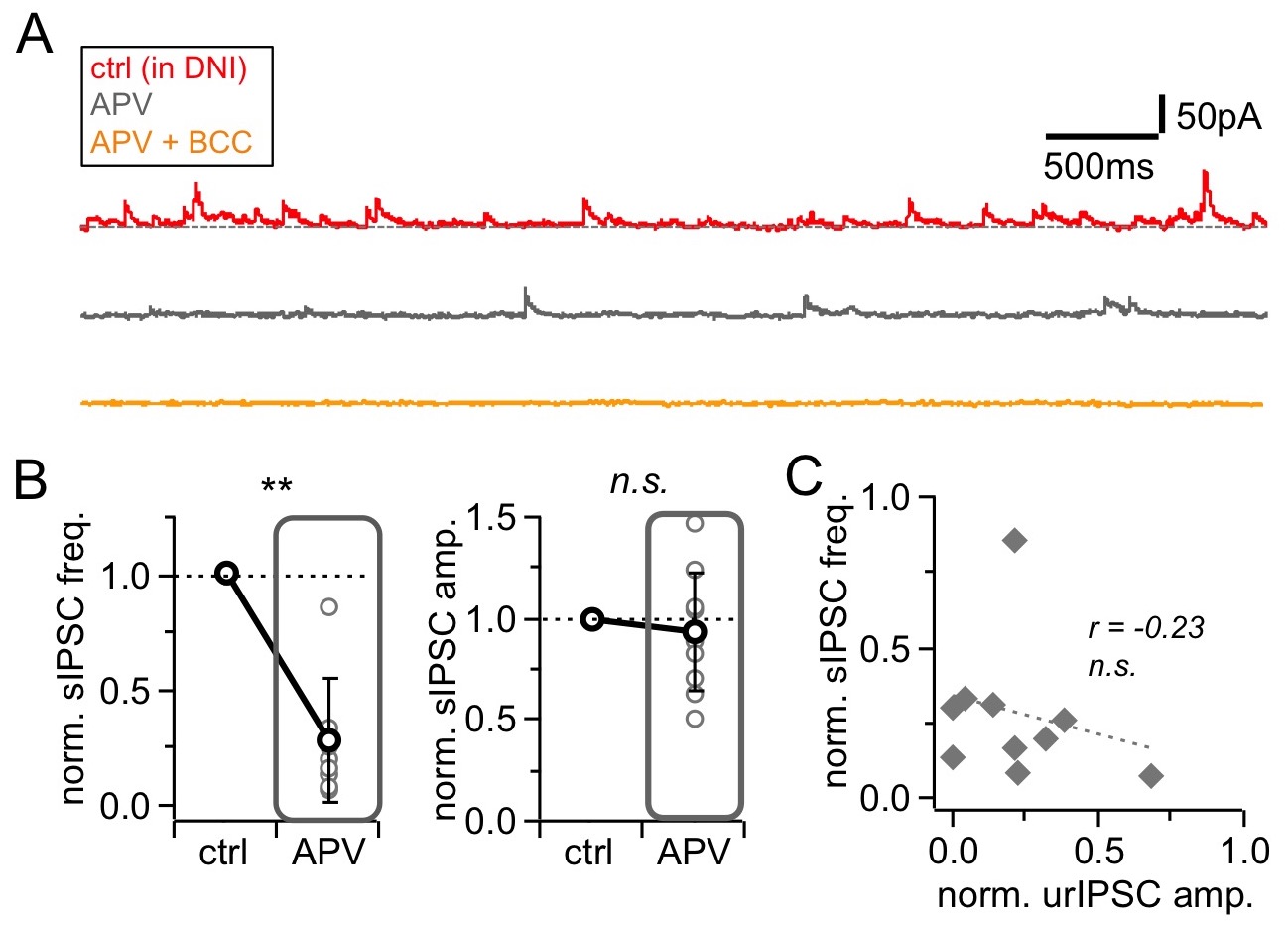

**Figure S4. Effects of NMDAR blockade by D-APV (APV, 25 µM) on spontaneous activity.**

(**A**) Representative experiment showing voltage-clamp recordings (+10 mV) from a mitral cell in the presence of DNI (control, top trace) upon addition of APV (middle trace) and further wash-in of BCC (50 µM; bottom trace).

(**B**) sIPSCs in control and in presence of APV (n = 10 MCs). Left: Frequency. Right: Amplitude. sIPSCs were abolished in BCC (analysis not shown) (**C**) Linear regression between the APV effect on sIPSC frequency versus urIPSC amplitude (n = 10 MCs). No significant correlation.
